# Novel bioinformatics approach to investigate quantitative phenotype-genotype associations in neuroimaging studies

**DOI:** 10.1101/015065

**Authors:** Sejal Patel, Min Tae M. Park, The Alzheimer’s Disease Neuroimaging Initiative, M. Mallar Chakravarty, Jo Knight

**Author notes:** **Correspondence**: Sejal Patel, Campbell Family Mental Health Research Institute, Centre for Addiction and Mental Health, 250 College Street, Toronto, ON, M5T 1R8, Canada. M. Mallar Chakravarty and Jo Knight are co-senior authors. Data used in preparation of this article were obtained from the Alzheimer’s Disease Neuroimaging Initiative (ADNI) database (adni.loni.usc.edu). As such, the investigators within the ADNI contributed to the design and implementation of ADNI and/or provided data but did not participate in analysis or writing of this report. A complete listing of ADNI investigators can be found at: http://adni.loni.usc.edu/wp-content/uploads/how_to_apply/ADNI_Acknowledgement_List.pdf.

## Abstract

Imaging genetics is an emerging field in which the association between genes and neuroimaging-based quantitative phenotypes are used to explore the functional role of genes in neuroanatomy and neurophysiology in the context of healthy function and neuropsychiatric disorders. The main obstacle for researchers in the field is the high dimensionality of the data in both the imaging phenotypes and the genetic variants commonly typed. In this article, we develop a novel method that utilizes Gene Ontology, an online database, to select and prioritize certain genes, employing a stratified false discovery rate (sFDR) approach to investigate their associations with imaging phenotypes. sFDR has the potential to increase power in genome wide association studies (GWAS), and is quickly gaining traction as a method for multiple testing correction. Our novel approach addresses both the pressing need in genetic research to move beyond candidate gene studies, while not being overburdened with a loss of power due to multiple testing. As an example of our methodology, we perform a GWAS of hippocampal volume using the Alzheimer’s Disease Neuroimaging Initiative sample.

## 1. Introduction

Imaging genetics is a burgeoning field that seeks to understand the association of neuroimaging-based phenotypes, such as structural, functional (Thompson *et al*., 2010) and diffusion imaging-based metrics, (Patel *et al*., 2010) with genetic variations. Candidate gene studies were initially the method of choice for understanding gene function in humans, and successfully identified genes involved in Mendelian diseases; however, such studies have had less success for complex genetic disorders, with many novel findings failing to replicate in further studies (Hirschhorn *et al*., 2002). Reasons for such failures include a lack of power to identify the small effect sizes typically involved in complex traits, as well as a lack of knowledge about which genes are appropriate to study (Tabor *et al*., 2002; Ioannidis, 2005). Around 2007, genome-wide association studies (GWAS) began to make inroads as an efficient method for identifying variants associated with complex disease. In this approach, approximately one million single nucleotide polymorphisms (SNPs) across the whole genome are interrogated simultaneously, hypothesis-free (Wellcome_Trust_Case_Control_Consortium, 2007). However, due to the large burden of multiple testing correction in a GWAS, a p-value of 5×10^−8^ or less, roughly equivalent to a p = 0.05 after Bonferroni correction for half a million independent variants, is generally required for a SNP to be recognized as significantly associated with a trait (Dudbridge and Gusnanto, 2008). Given the polygenic nature of complex traits and low effect sizes associated with these traits, large sample sizes are required to achieve adequate statistical power. Recently, a large imaging genetics study named ENIGMA (Enhancing NeuroImaging Genetics through Meta-Analysis) was undertaken, in which 21,000 subjects were included in a GWAS in order to identify genetic variants with association to hippocampal volume (Stein *et al*., 2012). While this study was a landmark demonstration for the use of imaging genetics techniques to investigate brain structures, it is not plausible for individual investigators to obtain such large sample sizes for their studies.

Various approaches have been described to reduce the multiple testing burden for large scale GWAS. One such approach is to control for the false discovery rate (FDR), rather than the family-wise error rate (FWER) (Benjamini and Hochberg, 1995). Where the family-wise error rate identifies the probability of one type 1 error from the total tested hypotheses, FDR calculates the proportion of expected type 1 errors. Stratified false discovery rate (sFDR) is an extension of the FDR control approach, where the false discovery rate is controlled in distinct subsets (strata) of the data, one or more of which are believed to have a higher prior probability of being associated with the trait of interest. Strata are defined based on prior information such as linkage analysis, candidate gene studies, or biological pathways (Sun *et al*., 2006; Sun *et al*., 2012). An example of this approach by Sun et al. (2012) investigates the susceptibility to meconium ileus (severe intestinal obstruction) in individuals with cystic fibrosis by prioritizing a set of genes involved in the apical plasma membrane. In this article, Sun et al. (2012) selected strata defined by Gene Ontology (GO) terms. GO is a biomedical ontology database, which contains structured vocabulary terms known as GO terms designed to describe protein function (Ashburner *et al*., 2000). In more complex traits, this approach may not be refined enough for the proper stratification of data in sFDR. In the case of Alzheimer’s disease (AD), for example, a vast number of SNPs would be selected to have prior association, and this may reduce the value of stratification. We present a novel method that employs information from previous studies alongside GO in order to create a refined list of relevant genes to be analyzed in the sFDR framework.

We demonstrate our method’s efficacy by investigating the association between genetic variants and hippocampal volume in the Alzheimer’s Disease Neuroimaging Initiative (ADNI1) dataset; sFDR is performed to stratify genes of particular interest to AD. The dominant symptom of AD is dementia, where memory, reasoning, and thinking are all impaired. The hippocampus plays a key role in cognitive functioning, influencing processes such as learning and the ability to make new memories (Braskie *et al*., 2013). Further, in considering the neurodegeneration of medial temporal lobe structures, the changes in hippocampal structure are considered to be one of the strongest quantitative phenotypes associated with AD and can often be used to predict cognitive decline in AD patients (Braskie *et al*., 2013).

This article presents a novel, systematic method to determine the optimal stratification of SNPs for sFDR analysis. We employed GO alongside previous GWAS findings, and applied our method to the ADNI1 dataset. Our method reduces the multiple testing correction burden with the potential to discover novel biomarkers in imaging genetics. Useful not only for new genetic studies, our tool is highly applicable to mining already existing GWAS data and improving the integration of publically available bioinformatics resources such as GO with imaging genetics studies.

## 2. Materials and Methods

### 2.1 Selecting Priority List of Genes

Below, we detail how we assembled a list of priority genes, derived comprehensive gene networks based on these so-called “seed” genes, and pruned these networks appropriately. Our priority SNPs were selected from genes involved in biological systems associated with AD. Figure 1 outlines the entire process followed, including SNP selection (Figure 1A) and the preparation and subsequent analysis of the genetic and imaging data (Figure 1B).

**Figure 1.**
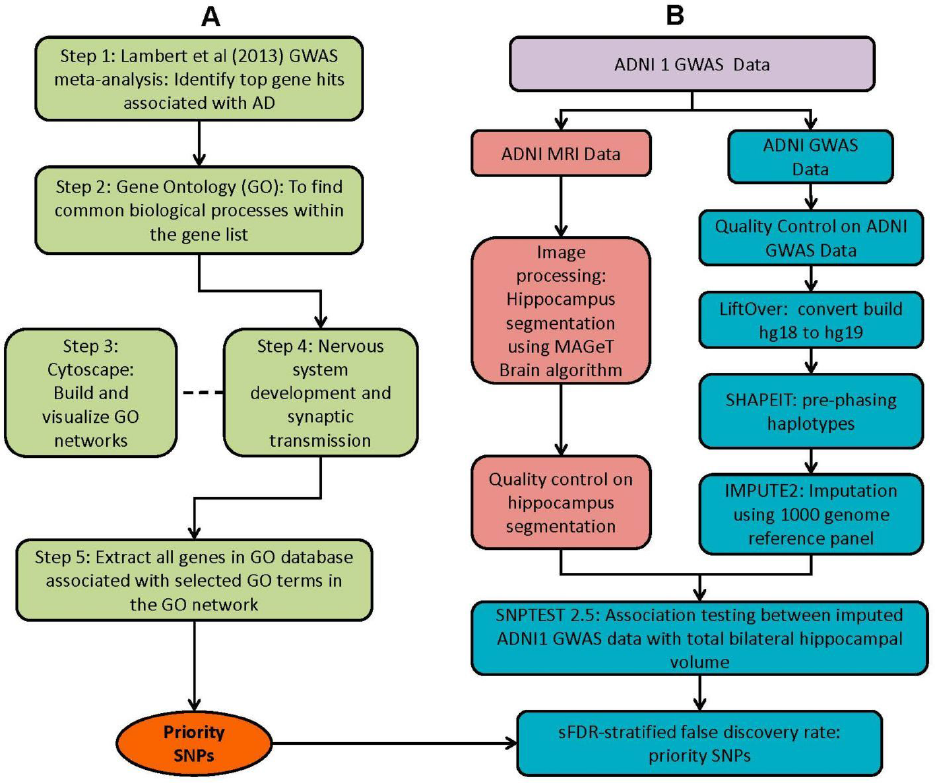
Method overview of both the selection of priority SNPs and association testing analysis between ADNI1 GWAS and imaging data. **(A)** Steps taken to select for priority SNPs. Gene hits from a meta-analysis by Lambert *et al*., (2013) were used as a starting point (Step 1) and GO was then used to identify common biological processes within the gene hits (Step 2). Cytoscape was used to build and visualize common biological process networks -- in this case the “nervous development and synaptic transmission” network was selected (Step 3 and Step 4). All genes from the selected GO terms in the network were extracted to form the priority list of SNPs. sFDR was then implemented with the priority SNPs. (**B)** Shows quality controls steps taken GWAS data and hippocampal imaging data. Association analysis was performed between imputed quality control (QC) GWAS data with QCed hippocampal segmentation.

**Step 1**: Twenty-one hits from a previous meta-analysis of AD GWAS signals were used as a starting point to identify top gene hits (Lambert *et al*., 2013). In addition we added amyloid precursor protein (*APP*) (Goate *et al*., 1991), Presenilin-1 (*PSEN1*) and Presenilin-2 (*PSEN2*) (Cruts *et al*., 1998) to our gene list based on association with higher risk of developing early onset of AD. Furthermore rare variants within these gene regions also increase the risk of late onset AD (Cruchaga *et al*., 2012).

**Step 2**: Gene Ontology (GO) (refer to Box 1) was used to group genes, and subsequently to derive common biological process networks using a three step process detailed below.

#### Box 1

##### Gene Ontology

Gene Ontology (http://geneontology.org/) is a publically available, free, ontology database that describes protein function (Ashburner *et al*., 2000). Gene products – proteins – are classified and grouped in three main ontologies: cellular components (CC) where the protein is located within subcellular compartments, molecular functions (MF) indicates the specific function of the gene is carried out in normal conditions and biological processes (BP) which describes the processes a protein is involved in (e.g.: neurogenesis). The ontology follows a hierarchical order and there are defined relationships between the GO terms. In the ontology structure, terms at the top represent general or broad concepts, whereas terms near the bottom represent more detailed processes. Therefore if a term has terms subordinate to it, it is referred to as a ‘parent’ term. Similarly, if a term has other terms superior to it, then it is referred to as a ‘child’ term. Both manual and automatic annotations of proteins are available in the GO database. Automatic annotations are inferred from electronic annotations and are not manually reviewed by a curator. In manual annotations, a curator reviews primary articles to generate annotations, and each annotation is based on experimental data referenced to a PubMed ID. The documentation for manual curation can be found at http://geneontology.org/page/annotation, and an example of annotations created by the authors can be found in the Alzheimer’s University of Toronto dataset at http://www.ebi.ac.uk/QuickGO/GAnnotation?source=Alzheimers_University_of_Toronto Quick GO (http://www.ebi.ac.uk/QuickGO/) is a web based tool used to extract data from the GO database.

##### Cytoscape

Cytoscape is an open source software platform visualization tool used to integrate data into complex networks of molecular interaction and biological pathways ((Saito *et al*., 2012), http://www.cytoscape.org/). See Figure 4 as an example of a biological network.

Firstly, the biological process (BP) ontology dataset within GO was examined using Quick GO (refer to Box 1) in order to identify all BP terms associated with the genes under investigation, hereafter called originally selected GO terms (OGO terms). No restrictions were given on the type of evidence codes used for the annotation of the OGO terms. Secondly, similar OGO terms were grouped together to form key BP domain categories. Specifically, they were grouped based on common parent terms, which were higher up in the hierarchal ontology. Thirdly, common biological processes were identified based on the frequency of occurrence of previously associated genes. Only OGO terms associated with these common processes are carried forward to the next step.

Based on the outcome of the three steps outlined above, we grouped all child terms that derived from parent terms in the domains of synaptic function, neuroanatomical structure development, and neurogenesis. These three parent terms were found to be under the common network of “nervous system development and synaptic transmission”, which was identified as a common biological process. GO terms that fell under the network “nervous system development and synaptic transmission” were the fourth child term from the BP parent GO term. Refer to Box 1 under Gene Ontology section for child and parent terminology. In order to benchmark our approach in the selection of common biological processes, INRICH (Lee *et al*., 2012) was used as an alternative, objective, method to derive the common biological process domains. However, no significant results were identified to take forward to sFDR. The INRICH process is defined in the supplementary text.

**Step 3**: Cytoscape 2.8 (refer to Box 1) was used to visualize the biological process network “nervous system development and synaptic transmission”, and parent GO terms from the OGO terms were extracted to contextualize this network. As expected, the networks were overly complex and contained much extraneous information. To remedy this, an algorithm was developed to effectively reduce redundant data in order to create an effectively “pruned” network. This is accomplished by using building and pruning techniques based on the relationships of OGO terms. Figures 2–4 demonstrate different stages of this algorithm with the OGO terms in green boxes. Figure 2 shows a subsection of GO terms in the complete “nervous system development and synaptic transmission” network before pruning of the data. Figures 3A to 3D display how specific criteria were used to remove non-targeted GO terms. Figure 4 shows the final pruned data of the nervous system development and synaptic transmission network.

**Figure 2.**
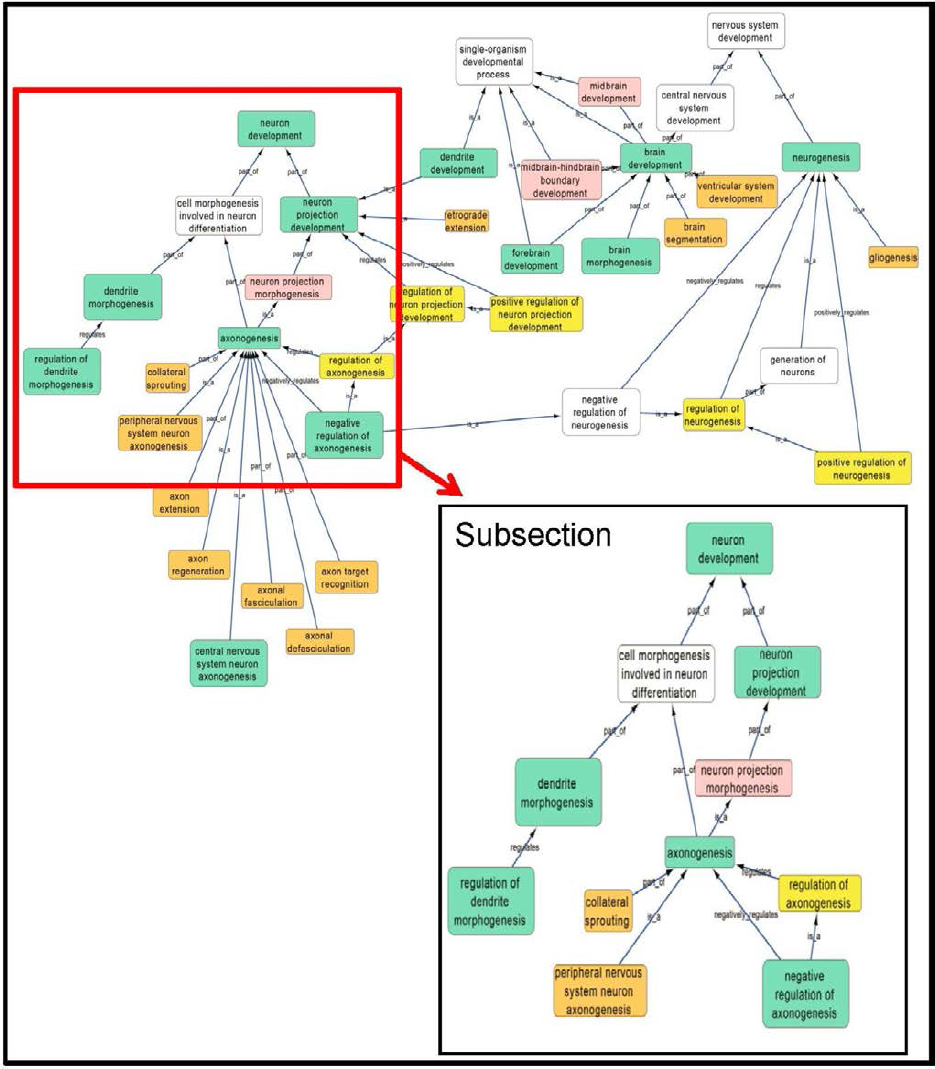
Sample of initial network with selected GO terms before pruning. A subsection is selected to show how the criteria was used to prune the complex GO network. Pruning steps are shown in Figure 3.

**Figure 3.**
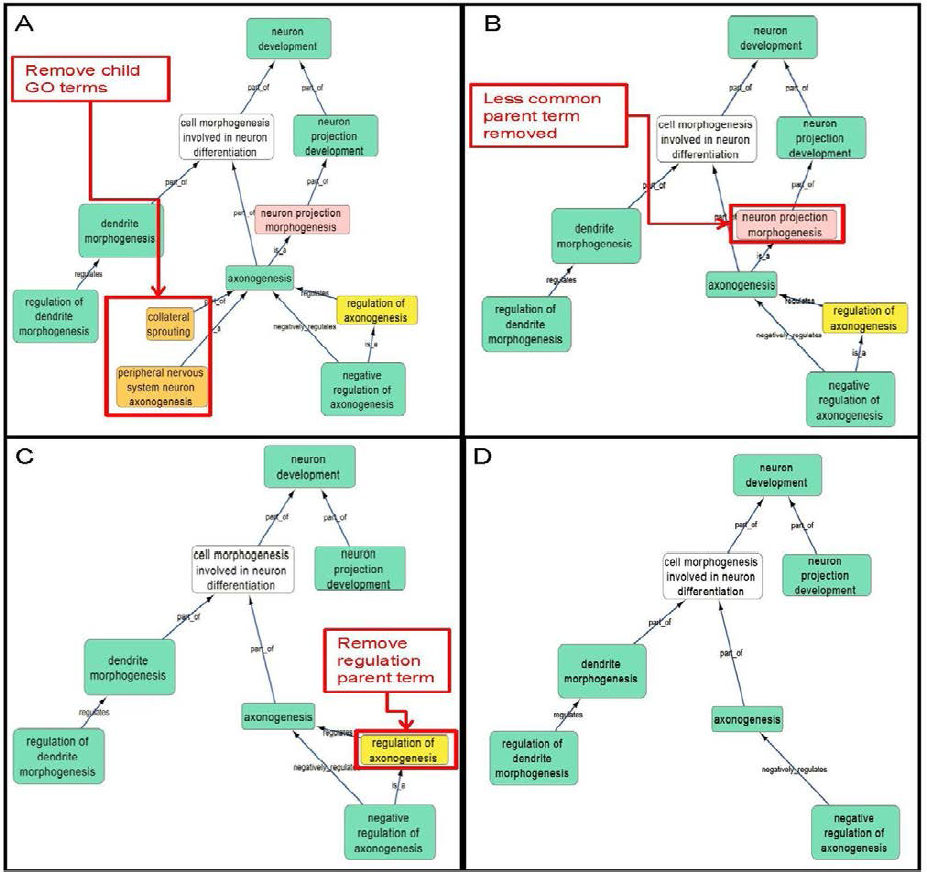
Criteria used to prune a complex network. Green box: Selected GO terms that are associated with a gene identified from Lambert *et al*., (2013). Orange box: Child terms of selected GO terms. Pink box: Less common parent term only associated with one selected parent GO term. Yellow box: Regulation GO terms that do not specify positive or negative regulation. (**A)** Child terms of selected GO terms were removed. (**B)** A less common parent GO term (neuron projection morphogenesis) which has one selected child GO term (‘axongenesis’) is removed because ‘Cell morphogenesis involved in neuron differentiation’ is a parent term for both selected GO terms ‘axonogenesis’ and ‘dendrite morphogenesis’. (**C)** Regulation terms that does not specify the type of regulation is removed because selected GO term ‘negative regulation of axonogensis” is more descriptive than the parent GO term ‘regulation of axongenesis’. **(D)** A sample of a pruned network after following the criteria in Figure 3A –3C.

The following criteria were used to select the child and parent GO terms.

a. When extracting the ontology of the OGO terms using Cytoscape, child terms are automatically selected. Therefore, to simplify the ontology networks, child terms were removed. Orange terms in Figure 3A represent extra child terms of the OGO terms, which are not needed in the network. For example the GO term ‘axonogenesis’ has two child terms ‘collateral sprouting’ and ‘peripheral nervous system neuron axonogenesis’. These are not necessary because the genes from step 1 have not been associated with these GO terms. (Figure 3A)
b. If more than one parent term is identified for an OGO term, then a common parent term, which is shared by most of the OGO terms, is chosen. As an example, the term ‘axonogenesis’ has two parent terms, namely, ‘neuron projection morphogenesis’ and ‘cell morphogenesis involved in neuron differentiation’. In Figure 3B the term ‘neuron projection morphogenesis’, displayed in a pink box, is removed because the alternate parent term, ‘cell morphogenesis involved in neuron differentiation’, is a parent term to both the selected GO terms ‘dendrite morphogenesis’ and ‘axonogenesis’.
c. A positive or negative regulation child term will have two types of parents. As an example, we will investigate the term ‘negative regulation of axonogenesis’. The first parent will be the term it regulates (‘axongenesis’) and the second parent would likely be a term that has ‘regulation’ as a key word in the term name, for example, ‘regulation of axonogenesis’ could be a candidate. Therefore the parent term that is regulated was selected, in this case the term ‘axonogenesis’, and the parent term that regulates a biological process but does not specify positive or negative regulation (‘regulation of axonogenesis’) is removed – shown in a yellow box -- because the child term will be more specific in terms of explaining how it is regulating the parent term (eg. negative regulation of axonogenesis), Figure 3C.

**Step 4**: Quick GO was used to extract all the genes that are associated to the OGO terms (as displayed in Figure 4 in green boxes) in the pruned “nervous system development and synaptic transmission” network. SNPs from these genes were extracted from the ADNI1 dataset using a reference file containing the start and end positions of the transcribed gene portion according to the Homo sapiens build 37 protein and coding genes from National Center for Biotechnology Information (NCBI). This list of SNPs formed the priority list for sFDR.

**Figure 4.**
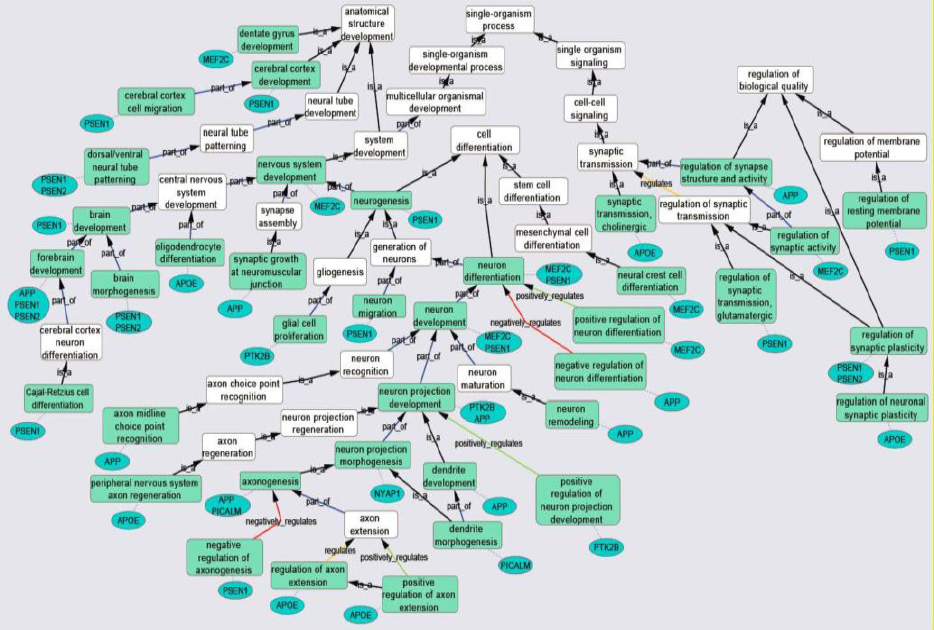
Gene Ontology (GO) biological process network of the “nervous system development and synaptic transmission” in association with AD. Green boxes are GO terms that are associated with the specific genes (blue ovals) connected by purple dotted line. White boxes are intermediate parent GO terms related to the selected GO terms (green boxes). Black arrows represent ‘is_a’ relationship between the GO terms and its parent term; blue arrows shows a ‘part _of’ relationship; orange arrows, a ‘regulation’ relationship; green arrows, a ‘positive_regulation’ relationship and red arrows, a negative_regulation’ relationship.

### 2.2 ADNI Imaging Data

#### 2.2.1 ADNI Data

GWAS data and magnetic resonance imaging (MRI) neuroimaging data was obtained from the Alzheimer’s Disease Neuroimaging Initiative (ADNI). Established in 2003 to facilitate the development of methods for biomarker investigation in order to enable detection of Alzheimer’s disease (AD) at earlier stages, ADNI is a partnership between the National Institute on Aging, the National Institute of Biomedical Imaging and Bioengineering, the Food and Drug Administration, private pharmaceutical companies, and nonprofit organizations (http://adni.loni.usc.edu/; Michael W. Weiner, Principal Investigator). The ADNI database contains different information including neuroimaging, clinical, and genome-wide SNPs data. According to the ADNI protocol, subjects are diagnosed as cognitively normal (CN), mild cognitive impairment (MCI), or Alzheimer’s disease (AD), based on the severity of their condition, and are recruited from Canada and the United States.

We used the ADNI1 dataset “ADNI1: Complete 1Yr 1.5T” (Wyman *et al*., 2013). 1.5T scanners (General Electric Healthcare, Philips Medical System or Siemens Medical Solutions) were used with the protocol described by (Jack *et al*., 2008). Before quality control (QC), 817 Caucasian 1.5T MRI subject scans were obtained from the ADNI1 database. Of the 817 subjects, 757 had GWAS data and 662 passed quality control. Figure 1B shows the overall steps taken to process the ADNI 1 MRI and GWAS data.

#### 2.2.2 Hippocampal Segmentation

Hippocampal segmentation was carried out in all 662 samples with GWAS data, using a modified multi-atlas algorithm known as the Multiple Automatically Generated Templates (MAGeT-Brain) algorithm (Chakravarty *et al*., 2013; Pipitone *et al*., 2014). The MAGeT Brain algorithm overcomes the limitations of model-based segmentation techniques, and avoids the requirement for larger atlas libraries typically required in more traditional multi-atlas segmentation strategies (Heckemann *et al*., 2006; Collins and Pruessner, 2010) by bootstrapping the segmentation procedure using data from the participants being analyzed. The segmentation procedure consists of three steps. First, five high-resolution MRI atlases developed by our group were used as inputs (Winterburn *et al*., 2013) and are used to automatically generate a “template library” based on a subset of the ADNI1 dataset using a model based segmentation procedure. For the purposes of this work we used a subset of subjects consisting of 7 AD, 7 MCI and 7 CN subjects evenly distributed across an age range of 58-90 to model the anatomical variability across the ADNI1 dataset. Model-based segmentation is used to segment each of the subjects in the template library leading to a total of 5 candidate segmentations per subject. The next step proceeds much like a regular multi-atlas segmentation strategy, where each subject is nonlinearly matched to each of the subjects in the template library, yielding 105 (5 atlases × 21 templates) candidate segmentations for each subject. The last step is a voxel voting technique where a label at each voxel that is most frequently occurring is used for the final segmentation (Collins and Pruessner, 2010). All resultant segmentations were manually inspected by an expert rater and only those segmentations passing quality control were used in the analysis. Images not successfully segmented by the MAGeT Brain algorithm were segmented manually for use. All input atlases (http://cobralab.ca/atlases/Hippocampus.html) and source code for MAGeT-Brain are freely available online (https://github.com/CobraLab/MAGeTbrain). Nonlinear transformations were estimated using the ANTs algorithm (Avants *et al*., 2008) and image processing steps were carried out using the MINC toolbox (http://www.bic.mni.mcgill.ca/ServicesSoftware/ServicesSoftwareMincToolKit).

### 2.3 ADNI1 Genetic Data

#### 2.3.1 Genetic Data Quality Control

Quality control (QC) was performed on the ADNI 1 GWAS data (N=757) using PLINK (version 1.07, http://pngu.mgh.harvard.edu/∼purcell/plink/ (Purcell *et al*., 2007)). In addition R (http://www.r-project.org/) was used to visualize the results. Individuals with discordant sex information, high level of missing data (> 2%) and heterozygosity rates greater than 3 standard deviations from the mean were removed from the sample. One of each pair of individuals displaying a high level of pair-wise identity by descent (IBD > 0.185) were also removed. In addition, SNPs with minor allele frequency (MAF) <1% and Hardy-Weinberg equilibrium (p < 1×10^−7^) were removed. After QC, 662 individuals remained in the analysis set. Multidimensional scaling (MDS) was performed in PLINK using HapMap3 (Altshuler *et al*., 2010) as a reference panel. When the population is compared with the CEU (CEPH - Utah residents with ancestry from northern and western Europe), YRI (Yoruba in Ibadan, Nigeria), JPT (Japanese in Tokyo, Japan), TSI (Tuscans in Italy) and CHB (Han Chinese in Beijing, China) ancestry, the sample clustered around CEU and TSI sample. MDS was subsequently carried out with the ADNI1, CEU, TSI and Jewish ancestry samples and aligned completely with the later three samples (Supplementary Figure S1). The Jewish ancestry sample was made available by Mark Silverberg.

#### 2.3.2 Data Preparation, Pre-Phasing and Imputation

The GWAS data was based on UCSC, (University of California, Santa Cruz) build 36 reference (Lander *et al*., 2001), and the liftover tool available from the NCBI (http://genome.ucsc.edu/cgi-bin/hgLiftOver) was used to convert each SNP location to build 37. SHAPEIT 2.0 ((Delaneau *et al*., 2012), https://mathgen.stats.ox.ac.uk/genetics_software/shapeit/shapeit.html) was used to pre-phase the haplotypes of the GWAS data after QC. Imputation was performed on the pre-phased data using Impute2 ((Marchini *et al*., 2007), https://mathgen.stats.ox.ac.uk/impute/impute_v2.html) for the autosomal chromosomes with the 1000 Genome (March 2012) data as a reference. SNPs with info values of equal and greater than 0.5 and MAF > 0.05 were retained for analysis.

#### 2.3.3 Association of Hippocampal Volume with GWAS Data

SNPTEST 2.5 ((Marchini *et al*., 2007), https://mathgen.stats.ox.ac.uk/genetics_software/snptest/snptest.html) was used to examine associations between hippocampal volumes with both imputed and genotyped SNPs. Covariates used in the analysis were gender, age, first dimension from MDS to control for population structure, baseline diagnoses (CN, MCI, or AD), APOE status because *APOE* e4 carriers have a higher risk of developing AD (Farrer *et al*., 1997) and intracranial volume to correct for variation in brain sizes within individuals in the sample. Three phenotypes were investigated: left hippocampal volume, right hippocampal volume, and mean (of the left and right) hippocampal volume. Frequentist association testing was undertaken for each phenotype, with a ‘method’ option in place to control for genotype uncertainty in the association test.

#### 2.4 Stratification of SNPs

Fixed FDR strategies are used to control FDR in a group of tests. In sFDR, SNP p-values from the association analysis are grouped into distinct strata, one or more of which are believed to have a higher prior probability of being associated with the trait of interest (Sun *et al*., 2006). The association p-values of each SNP are transformed to q-values and FDR is controlled separately within each strata. To control the FDR at a given level – 5% in this analysis – the null hypothesis is rejected when tests have a q-value equal to or less than the specified threshold (0.05). This method increases the power to identify true associations if one of the strata is enriched with associated variants. When the strata aren’t enriched, the method is still robust. Two SNP strata were formed in our data. All SNPs in the genes associated to the OGO terms (Figure 4) from the pruned “nervous system development and synaptic transmission” network formed one, high priority, strata (249,001 SNPs), and all the remaining SNPs formed the other (5,457,557 SNPs) in our non-priority stratum. Association p-values from SNPTEST were merged with each corresponding SNPs in each strata (priority and non-priority list) for sFDR. A Perl script was used to analyze priority and non-priority SNPs (http://www.utstat.toronto.edu/sun/Software/SFDR/).

## 3. Results

### 3.1 SNP Selection

**Step 1**: From the 21 loci identified in Lambert *et al*., (2013) in association with AD, 10 were already known through previous GWAS and 11 novel loci were found (Table 1). *APP*, *PSEN1* and *PSEN2* were also added to the gene list.

**Table 1.**
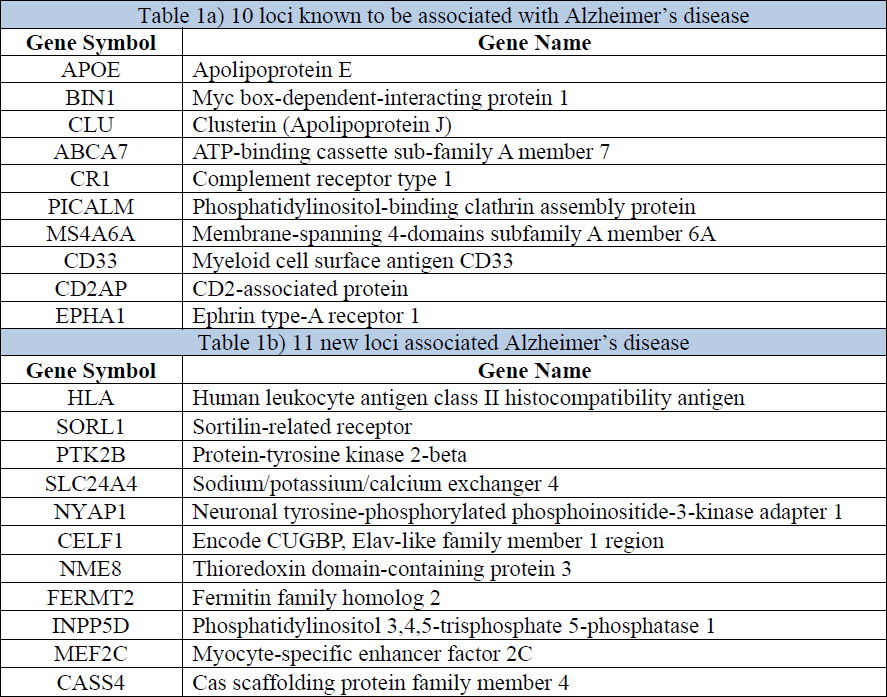
Top genes associated with AD from the Lambert *et al*., (2013) meta-analysis.

| Table 1a) 10 loci known to be associated with Alzheimer's disease |  |
| --- | --- |
| Gene Symbol | Gene Name |
| APOE | Apolipoprotein E |
| BIN1 | Myc box-dependent-interacting protein 1 |
| CLU | Clusterin (Apolipoprotein J) |
| ABCA7 | ATP-binding cassette sub-family A member 7 |
| CR1 | Complement receptor type 1 |
| PICALM | Phosphatidylinositol-binding clathrin assembly protein |
| MS4A6A | Membrane-spanning 4-domains subfamily A member 6A |
| CD33 | Myeloid cell surface antigen CD33 |
| CD2AP | CD2-associated protein |
| EPHA1 | Ephrin type-A receptor 1 |
| Table 1b) 11 new loci associated Alzheimer's disease |  |
| Gene Symbol | Gene Name |
| HLA | Human leukocyte antigen class II histocompatibility antigen |
| SORL1 | Sortilin-related receptor |
| PTK2B | Protein-tyrosine kinase 2-beta |
| SLC24A4 | Sodium/potassium/calcium exchanger 4 |
| NYAP1 | Neuronal tyrosine-phosphorylated phosphoinositide-3-kinase adapter 1 |
| CELF1 | Encode CUGBP, Elav-like family member 1 region |
| NME8 | Thioredoxin domain-containing protein 3 |
| FERMT2 | Fermitin family homolog 2 |
| INPP5D | Phosphatidylinositol 3,4,5-trisphosphate 5-phosphatase 1 |
| MEF2C | Myocyte-specific enhancer factor 2C |
| CASS4 | Cas scaffolding protein family member 4 |

**Step 2**: Common biological processes within the gene list were identified using GO. INRICH was used as an alternative objective method, but significant results were not found. Regardless, results from INRICH may be found in the supplementary section. The GO database was accessed on January 27^th^ 2014. In the GO database all genes from the list had BP GO terms annotated to them except the gene Membrane-spanning 4-domains subfamily A member 6A (*MS4A6A*). Table 2 shows the common BP domains associated with the 21 genes. In this study we focused on the “nervous system development and synaptic transmission” network (Figure 4), which included many genes from our original list. The network can be broken down into sub-domains with key GO terms in the areas of synaptic function, neuroanatomical structure development, and neurogenesis. For example, in the domain “neuroanatomical structure development”, myocyte-specific enhancer factor 2C (*MEF2C*) has been associated with GO terms ‘denate gyrus development’ and ‘nervous system development’.

**Table 2.** Common GO Biological Process domains of gene hits from the Lambert *et al*., (2013) meta-analysis.

| Gene | Protein ID | Vesicle-mediated transport and Endocytosis | Steroid and cholesterol metabolic process | Immune system process | Cell membrane processes and Cell migration | Nervous system development and Synaptic transmission | Regulation of calcium-mediated signaling |
| --- | --- | --- | --- | --- | --- | --- | --- |
| DRB5 | Q30154 |  |  | X |  |  |  |
| SORL1 | Q92673 | X | X |  | X |  |  |
| PTK2B | Q14289 |  |  | X | X | X | X |
| SLC24A4 | Q8NFF2 | X |  |  |  |  | X |
| NYAP1 | Q6ZVC0 |  |  |  |  | X |  |
| MADD | Q8WXG6 |  |  |  | X |  |  |
| NME8 | Q8N427 |  |  |  | X |  |  |
| FERMT2 | Q96AC1 | X |  |  | X |  |  |
| INPP5D | Q92835 |  |  | X |  |  |  |
| MEF2C | Q06413 |  |  |  |  | X |  |
| CASS4 | Q9NQ75 |  |  |  |  |  |  |
| APOE | P02649 | X | X |  | X | X | X |
| BIN1 | O00499 | X |  |  | X |  |  |
| CLU | P10909 | X | X | X |  |  |  |
| ABCA7 | Q8IZY2 | X | X | X |  |  |  |
| CR1 | P17927 |  |  | X |  |  |  |
| PICALM | Q13492 | X |  | X |  | X |  |
| MS4A6A | Q9H2W1 |  |  |  |  |  |  |
| CD33 | P20138 |  |  | X | X |  |  |
| CD2AP | Q9Y5K6 |  |  |  | X |  |  |
| EPHA1 | P21709 |  |  | X | X |  |  |

**Step 3**: Cytoscape visualization of the nervous system synaptic transmission network is shown in Figure 4.

**Step 4**: The list of genes associated with the OGO terms from the pruned nervous system and synaptic transmission network included 1249 genes, after removal of all non-autosomal genes 1146 genes remained and formed our stratum for sFDR. Supplementary Table S1 shows a list of all priority genes with chromosome number, start and end position and gene symbol. Furthermore Supplementary Table S2 contains all 249,001 SNPs from 1146 genes used for sFDR.

### 3.2 Quality Control of Imaging and GWAS data

After quality control (QC) of automatic hippocampal segmentations, 9 segmentations out of 662 subjects failed which were corrected though manual segmentation. For the ADNI1 GWAS data, the sample initially consisted of 757 individuals, and after QC the sample was reduced to 662 subjects. The number of SNPs in the GWAS data after QC was 529,623 from 620,901 original variants, of which 517,064 SNPs were on autosomal chromosomes. After imputation of the GWAS, data the number of SNPs typed increased to 17,418,272. After QC of imputed SNPs, 5,706,558 SNPs were used for the association analysis with mean hippocampal volume.

### 3.3 Association Testing with Hippocampal Volume

P-values from association testing between the SNPs and mean hippocampal volume did not result in any GWAS significant findings after correction for multiple testing (Figure 5). Some, however, approached significance (Table 3, SNPs with uncorrected p-values). For example rs72909661 in gene region Stearoyl-CoA desaturase 5 (*SCD5)* neared GWAS significance with an uncorrected p= 8.97×10^−7^. The top 10 SNPs found within gene regions were: Autism susceptibility gene 2 protein (*AUTS2*; rs2158616; p= 1.16×10^−6^), Transmembrane protein - family with sequence similarity 155 member A (*FAM155A*; rs1033880; p=4.42×10^−6^) and long non-coding RNAs (LOC440173; rs11791915; p=1.76×10^−6^). Testing with left and right hippocampal volumes as response variables resulted in no GWAS significant findings, as displayed in the supplemental materials.

**Figure 5.**
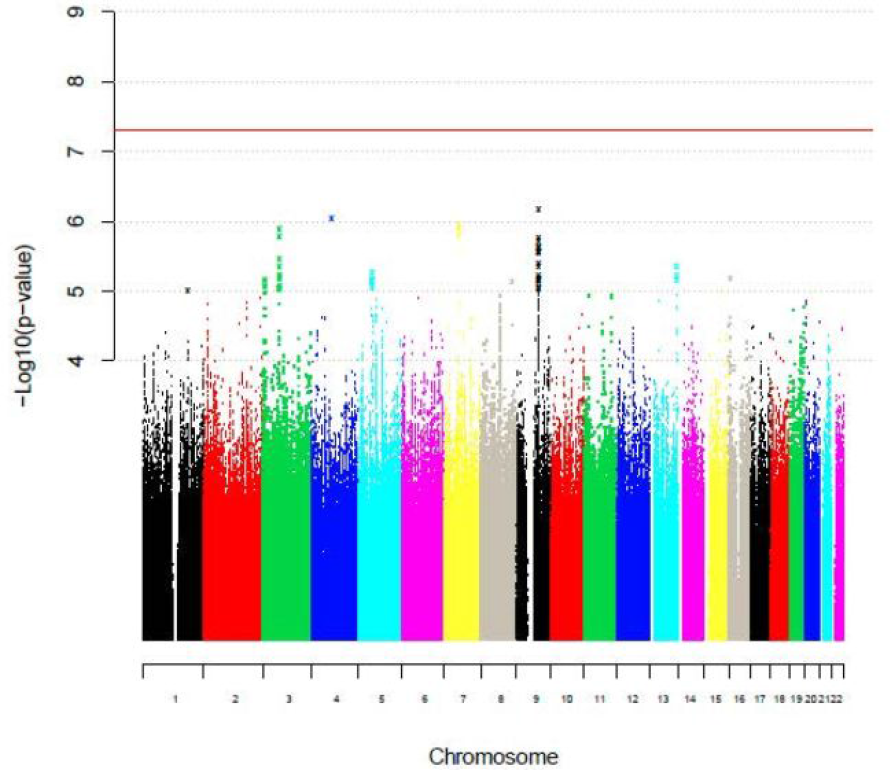
A Manhattan plot of imputed ADNI1 GWAS data. The x-axis represents the chromosomal location for each SNPs. The y-axis represents the log p-values of SNPs in association with AD. The red horizontal line represents the threshold for GWAS significant SNPs.

**Table 3.**
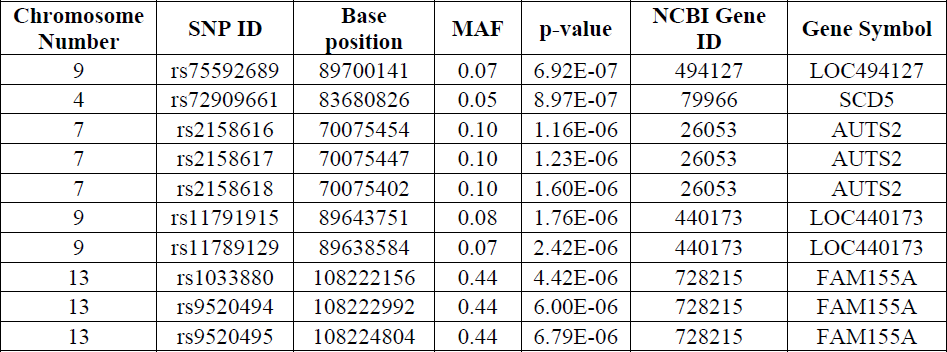
Top 10 most significant SNPs within gene regions from association testing of mean hippocampal volume. MAF represents minor allele frequency and p-value is the associated significance between the SNP and phenotype (mean hippocampal volume). Significant SNPs at a GWAS level are at p < 5×10^−8^.

| Chromosome Number | SNP ID | Base position | MAF | p-value | NCBI Gene ID | Gene Symbol |
| --- | --- | --- | --- | --- | --- | --- |
| 9 | rs75592689 | 89700141 | 0.07 | 6.92E-07 | 494127 | LOC494127 |
| 4 | rs72909661 | 83680826 | 0.05 | 8.97E-07 | 79966 | SCD5 |
| 7 | rs2158616 | 70075454 | 0.10 | 1.16E-06 | 26053 | AUTS2 |
| 7 | rs2158617 | 70075447 | 0.10 | 1.23E-06 | 26053 | AUTS2 |
| 7 | rs2158618 | 70075402 | 0.10 | 1.60E-06 | 26053 | AUTS2 |
| 9 | rs11791915 | 89643751 | 0.08 | 1.76E-06 | 440173 | LOC440173 |
| 9 | rs11789129 | 89638584 | 0.07 | 2.42E-06 | 440173 | LOC440173 |
| 13 | rs1033880 | 108222156 | 0.44 | 4.42E-06 | 728215 | FAM155A |
| 13 | rs9520494 | 108222992 | 0.44 | 6.00E-06 | 728215 | FAM155A |
| 13 | rs9520495 | 108224804 | 0.44 | 6.79E-06 | 728215 | FAM155A |

### 3.4 sFDR Results

In total there were 249,001 SNPs in our priority stratum and 5,457,557 SNPs in our non-priority stratum. No q-values from the priority list (nervous system development and synaptic transmission stratum) reached the 0.05 threshold. All of the top 97 ranked SNPs were found in our priority list, but these SNPs were not significant at a q-value of less than 0.05. In particular, SNPs in our priority list within gene regions: C2 calcium-dependent domain containing 3 (*CDCD3*), growth arrest-specific (*GAS7*), Semaphorin-3D (*SEMA3D*), Gap junction alpha-1 protein (*GJA1*) and Rap guanine nucleotide exchange factor 1 (*RAPGEF1*) were ranked in the top 10 but with an sFDR q-value of 0.71 (Table 4). Both the priority and non-priority stratum display uniform p-values; this indicates that there is no true association between the SNPs and mean hippocampal volume (Supplementary Figure S1 and S2).

**Table 4.** Top 10 most significant sFDR results for mean hippocampal volume. P-value is the associated significance between the SNP and phenotype (mean hippocampal volume). Significant SNPs at a GWAS level is p < 5×10^−8^. The sFDR q-value controls the false discovery rate; the q-value is the adjusted p-value. Significant q-value is set to 0.05. ‘Rank’ is the order of SNPs based on sFDR q-values from a total of 5,706,558 SNPs.

| Chromosome number | SNP ID | Base position | p-value | q-value | rank | NCBI Gene ID | Gene Symbol |
| --- | --- | --- | --- | --- | --- | --- | --- |
| 11 | rs12417424 | 73751481 | 2.94E-05 | 0.7113 | 1 | 26005 | C2CD3 |
| 17 | rs62064532 | 9850440 | 3.17E-05 | 0.7113 | 2 | 8522 | GAS7 |
| 11 | rs4944877 | 73753671 | 3.74E-05 | 0.7113 | 3 | 26005 | C2CD3 |
| 11 | rs58006161 | 73860874 | 4.05E-05 | 0.7113 | 4 | 26005 | C2CD3 |
| 11 | rs58719576 | 73860873 | 4.05E-05 | 0.7113 | 5 | 26005 | C2CD3 |
| 7 | rs17558985 | 84643942 | 4.19E-05 | 0.7113 | 6 | 223117 | SEMA3D |
| 17 | rs55753206 | 9856314 | 6.34E-05 | 0.7113 | 7 | 8522 | GAS7 |
| 6 | rs113413235 | 121762030 | 6.35E-05 | 0.7113 | 8 | 2697 | GJA1 |
| 9 | rs7469510 | 134559741 | 6.37E-05 | 0.7113 | 9 | 2889 | RAPGEF1 |

## 4. Discussion

In contrast to existing approaches, our novel method provides a systematic integration framework for previous knowledge with the GO database. Alternatives such as Aligator (Holmans *et al*., 2009) and INRICH both rely on the identification of over-represented GO categories among significant hits; we identify and adapt relevant categories and use sFDR to increase power while controlling for multiplicity.

With no individual variants reaching genome-wide significance, we suspect a lack of statistical power even while employing the sFDR framework. In the future, our method will be adapted and applied to different datasets, and specifically, alternate phenotypes of AD. The high concentration of amyloid plaques in subcortical structures in late stage AD and neurofibrillary tangles originating in the medial temporal lobe implicate these regions as potential targets for future analysis (Braak and Braak, 1991).

The process by which priority SNPs are selected for sFDR could be considered subjective and represents an area of active development for our algorithm. Often the associations in GWAS studies are designated to the most promising gene in the region from a biological standpoint, introducing bias in step 1. One approach to combat this phenomenon would be for the input list to include all genes within a high recombination region alongside the most significant hit. Selection of the common biological domains in step 2 is also subjective and could be replaced with a standard pathway approach such as INRICH or Aligator. We have performed pilot work using INRICH as outlined in the supplementary text. In this example no pathways were identified, preventing us from pursuing this avenue. This is likely due to a lack of power to identify relevant pathways.

Another area of active development relating to SNP selection revolves around growing and pruning the network of terms utilized. Specifically, the GO terms in the “nervous system development and synaptic transmission” domain are general, and apply towards whole brain structure; currently the ontology does not capture particular brain regions in details such as hippocampal development. For example, in our priority list, SNPs in gene region growth arrest-specific protein 7 which plays a role in neuronal development are mainly expressed in mature cerebellar Purkinje cells (Ju *et al*., 1998). As such, further pruning the GO network by focusing on BP GO terms annotated to genes specific to one brain region, such as the hippocampus, or neuronal cell type within such structures may be crucial. Biological processes associated with structural information have recently begun to be captured in GO (Huntley *et al*., 2014). Therefore, when genes are annotated to BP GO terms, additional information on where the biological process is occurring can be recorded. As a result, filtering the data and looking at BP GO terms occurring in neuro-anatomical cells in region of the hippocampus may help in further pruning the network.

Both automatic and manual curation was used to assign GO terms to the genes in question. Automatic curation is the result of machine learning algorithms, and the terms assigned tend to be much broader than the manually curated ones and adds a potential source of noise to our priority SNPs stratum. In the example analysis presented here the inclusion of these sub-optimal classifications was necessary due to the limited manual annotation of the loci observed in Lambert *et al*., yet we acknowledge the shortcomings of this approach and advise the prioritization of manually curated data GO data. To further address the issue we are in the process of manually curating the list of 21 loci associated with AD.

In conclusion, this article introduces the use of GO, an online database, as a novel method to efficiently prioritize data for sFDR multiple testing control. In particular we applied this method to a GWAS of hippocampal volume in the ADNI1 dataset. Though novel biomarkers were not identified, our method has the potential to improve the identification of genes in imaging-genetic studies.

## Acknowledgments

Computations were performed on the CAMH Specialized Computing Cluster. The SCC is funded by: The Canada Foundation for Innovation, Research Hospital Fund. Computations were performed on the gpc supercomputer at the SciNet HPC Consortium (Loken *et al*., 2010). SciNet is funded by: the Canada Foundation for Innovation under the auspices of Compute Canada; the Government of Ontario; Ontario Research Fund - Research Excellence; and the University of Toronto. MMC is funded by the National Sciences and Engineering Research Council, Canadian Institutes for Health, Weston Brain Institute, Michael J. Fox Foundation for Parkinson’s Research, Alzheimer’s Society, and Brain Canada. JK holds the Joanne Murphy Professorship in Behavioral Science and is also funded by the Canadian Institutes for Health. Sejal Patel is funded by The Cadsby Foundation. Thanks to Mark Silverberg for providing the Jewish ancestry sample. We would like to thank Chris Coles for his help with editing the manuscript.

ADNI Acknowledgments: Data collection and sharing for this project was funded by the Alzheimer’s Disease Neuroimaging Initiative (ADNI) (National Institutes of Health Grant U01 AG024904) and DOD ADNI (Department of Defense award number W81XWH-12-2-0012). ADNI is funded by the National Institute on Aging, the National Institute of Biomedical Imaging and Bioengineering, and through generous contributions from the following: Alzheimer’s Association; Alzheimer’s Drug Discovery Foundation; BioClinica, Inc.; Biogen Idec Inc.; Bristol-Myers Squibb Company; Eisai Inc.; Elan Pharmaceuticals, Inc.; Eli Lilly and Company; F. Hoffmann-La Roche Ltd. and its affiliated company Genentech, Inc.; GE Healthcare; Innogenetics, N.V.; IXICO Ltd.; Janssen Alzheimer Immunotherapy Research & Development, LLC.; Johnson & Johnson Pharmaceutical Research & Development LLC.; Medpace, Inc.; Merck & Co., Inc.; Meso Scale Diagnostics, LLC.; NeuroRx Research; Novartis Pharmaceuticals Corporation; Pfizer Inc.; Piramal Imaging; Servier; Synarc Inc.; and Takeda Pharmaceutical Company. The Canadian Institutes of Health Research is providing funds to support ADNI clinical sites in Canada. Private sector contributions are facilitated by the Foundation for the National Institutes of Health (www.fnih.org). The grantee organization is the Northern California Institute for Research and Education, and the study is coordinated by the Alzheimer’s Disease Cooperative Study at the University of California, San Diego. ADNI data are disseminated by the Laboratory for Neuro Imaging at the University of Southern California.

## Author Contributions

SP performed quality control and imputation on GWAS data, developed the Gene Ontology network and performed the association analysis between GWAS data and mean hippocampal volume. MTMP carried out automatic and manual hippocampal segmentation and quality control on the segmentation. SP, JK, MC conceptualized the design of the study. All authors wrote and edited the manuscript.

